# Establishment of baseline sensitivity of *Rhizoctonia solani* to thifluzamide in corn and its field application

**DOI:** 10.1101/402529

**Authors:** Dianlong Shang, Chentao Yao, Falin He, Xiao Sun, Shiang Sun, Haili Tan, Xiangdong Li, Jiwang Zhang, Xingyin Jiang

## Abstract

In recent years, banded leaf sheath blight in corn has become an important disease in corn that seriously affects quality and yield. This paper aims to evaluate the sensitivity of *Rhizoctonia solani* to thifluzamide in corn, to clarify the effect of seed coating using a thifluzamide suspension agent on safety and physiological indicators and to determine banded leaf sheath blight in corn control effectiveness in the field, thereby providing a basis for the application of thifluzamide suspension agent as a seed coating. In this study, the thifluzamide sensitivity of 102 strains of *R. solani* in corn in different regions of Shandong was determined using the mycelial growth rate method, and the average half-maximal effective concentration value (EC_50_) was 0.086±0.004 μg/mL. The sensitivity was consistent with a continuous and skewed normal distribution, and the sensitivity distribution frequency exhibited a continuous, unimodal curve, indicating that thifluzamide had strong inhibitory activity on the mycelial growth of *R. solani* in corn. The impacts of using a thifluzamide suspension agent for seed coating on safety and physiological indicators as well as the control effect in corn were evaluated by combining seed coating, an indoor pot test, and a field trial. The root activities under 24 g a.i. 100 kg^-1^ seed and 12 g a.i. 100 kg^-1^ seed were found to increase by 78.01% and 77.40%, respectively, compared with that under the blank control; the chlorophyll content of corn increased most significantly at a dosage of 24 g a.i. 100 kg^-1^, which was a 32.32% increase compared to the blank control. Thifluzamide (FS) could significantly increase the hundred-grain weight of corn and the per-plot yield. Among the examined dosages, 24 g a.i. 100 kg^-1^ seed had the most significant treatment effect, with the hundred-grain weight increasing by 12.47% and the yield rate increasing by 15.72% compared to the control in 2016, Simultaneously, the hundred-grain weight increasing by 13.44% and the yield rate increasing by 14.11% compared to the control in 2017. Three dosages of 24% thifluzamide (FS) increased the emergence rate and seedling growth of corn to varying extents. The field control effectiveness against banded leaf sheath blight in corn was best at the dosage of 24 g a.i. 100 kg^-1^ seed for seed dressing with thifluzamide (FS); in 2016 and 2017, the control effects in the small bell stage, large bell stage, tasseling and pollen-shedding stage, silking stage, milk-ripening stage, and wax-ripening stage were 100%, 66.73%, 52.8%, 67.81%, 68.48%, and 62.68% (2016), respectively, and 74.97%, 63.17%, 50.90%, 53.60%, 61.42%, and 55.88% (2017). These results indicated that thifluzamide had enormous potential for controlling banded leaf sheath blight in corn.

## Introduction

To promote the integrated control of air pollution to construct an ecological civilization in recent years, straw burning has been fully prohibited, while straw returning has been widely promoted in various places throughout China. However, due to improper treatment methods, straw returning has provided habitats for many soil-borne pathogens. As an important cereal crop in the global agricultural economy [1], corn is critical to increasing grain yield, but the incidence of banded leaf sheath blight in corn has been increasing annually, resulting in a decline in the quality and yield of corn and serious economic losses. Currently, farmers have a weak sense of prevention and control of banded leaf sheath blight in corn, and there is little use of control agents. Therefore, the development of safe, efficient agents for the prevention and treatment of this disease is urgently needed.

*Rhizoctonia* spp [2]. are destructive soilborne pathogens of many crops around the world that can utilize organic residues in the soil during the saprophytic period to survive as an aseptic mycelium (mycelium or sclerotia) [3,5]. Banded leaf sheath blight in corn is a soil-borne disease caused by infection by fungi in the soil habitat [6] such as *Rhizoctonia cerealis, Rhizoctonia solani*, and *Rhizoctonia zeae*. *Rhizoctonia solani* is a dominant pathogen in Shandong Province, China [7]. Its sexual stage is *Thanatephorus cucumeris*, and its main floras include AG-1-IA, AG-1-IB, AG-3, AG-5, AG-A, and AG-K [8,9,10]. The isolated strain of AG-1-IA readily causes banded leaf sheath blight in corn [11]. Disease incidence can span from the seedling period to the late growth period and be severe in the event of crop rotation [4,5]. The infection begins at the base of the leaf sheath, and peak damage occurs during the period from tasselling (VT) to grain filling. Initially, leaf sheaths have dark-green hygrophanous spots that gradually develop into cloud-shaped/wavy or irregular lesions from the bottom upward. The lesions are brown with the colour gradually becoming lighter from the outside to the inside; then, the lesions continue to expand and result in rotting of the leaf sheaths. In severe cases, stems become rotted and lodged/broken [12,13], and ears and grains become infested, causing insufficient grain filling, which seriously affects the quality and yield of corn[14].

At present, the methods for preventing and controlling banded leaf sheath blight in corn mainly include agricultural control, biological control, and chemical control, among which agricultural control has a limited effect and is time and labour consuming. Biological control has become an important area of research in plant protection in recent years. Tagele found that KNU17BI1 has great potential to control banded leaf sheath blight in corn caused by *R. solani* AG-1 (IA) [15], but the control effect is not ideal due to the limits of the growth environment. Hence, chemical control is still the most important prevention and control method in agricultural production. A previous study showed that the control effect of 25% triadimefon wettable powder (WP) could reach 44.17% when a 200-fold solution is applied for soil disinfection [16], and the control effect of 20% Jinggang mycin (AF) in fertilizer can exceed 80.1%. In addition, triazole fungicides, such as tebuconazole, have been used. Traditional control methods involve foliar spraying during the corn tasseling stage, which is limited by the height of the corn plants and is time-consuming and labourious. Thifluzamide is a thiazole amide fungicide that has both protection and treatment effects, and it can be used as a foliar spray or for soil treatment and can be quickly absorbed by plants. Thifluzamide is mainly used to prevent and control diseases caused by *Rhizoctonia* spp. of the phylum Basidiomycota [17,18].

Corn seed coating technology has also been widely used in corn planting. Through seed coating, the active ingredients of fungicides/pesticides are slowly released, which can, to some extent, enhance plant resistance and promote plant growth [19,20,21], thus having beneficial effects for corn [22]. In China, thifluzamide has achieved a good control effect as an agent against rice sheath blight. However, this effect has not been registered for corn, and no study on the control of banded leaf sheath blight in corn by seed dressing with thifluzamide has been reported. As a specific control agent of *Rhizoctonia* spp., investigating thifluzamide (FS) for the prevention and control of banded leaf sheath blight in corn is of great value. In this study, the baseline sensitivity of *R. solani* to thifluzamide was established in corn; the safety of thifluzamide coating was evaluated in corn, and the effects of thifluzamide on physiological and biochemical indicators of corn and its control of banded leaf sheath blight in corn were studied through pot and field fungicide tests to provide a basis for the application of a thifluzamide suspension agent for seed coating.

## Materials and methods

### Test materials

Test strains: In 2016-2017, diseased leaf sheaths, leaves, and stalks subjected to banded leaf sheath blight in corn were collected in 6 regions of Shandong, China: Tai’an (TA), Linyi (LY), Weifang (WF), Laiwu (LW), Rizhao (RZ), and Qingdao (QD). Upon isolation and purification, 102 strains of *R. solani* in corn were obtained. The sampling fields were never exposed to any thifluzamide or other SDHI. The identities of all isolates in the study were confirmed by morphology, phylogenetic analysis and pathogenicity testing. Isolates were kept for long-term storage in cryogenic tubes with 15% glycerol solution at –80°C. The test corn variety in this study was Zhengdan 958, (Henan Goldoctor Seed Co., Ltd., China). Test agents: The thifluzamide (96% TC; Shandong Kangqiao Bio-technology Co., Ltd.); the tebuconazole (94.7% TC; Shandong Weifang Runfeng Chemical Co., Ltd.); the thifluzamide (24% FS; made in the laboratory; Contains the following materials: FS3000, FS7PG, 2%XG, Deionized water, Magnesium aluminium silicate, White carbon black, LXC, D625, EP60P, Film former); and the 60 g/liter tebuconazole (FS) was provided by Bayer CropScience (China) Co., Ltd.

### Establishment of baseline sensitivity of *Rhizoctonia solani* to thifluzamide in corn

The mycelial growth rate method was used to determine the susceptibility of each of the 102 strains to thifluzamide, and a baseline sensitivity was established. Thifluzamide was dissolved with acetone and was prepared as a 500-μg/mL stock solution with 0.1% Tween-80 and sterilized deionized water. Using the stock solution for dilution, drug-containing PDA plates with thifluzamide concentrations of 1, 0.5, 0.25, 0.125, and 0.0625 μg/ml were prepared; a PDA plate with the same volume of sterilized water was used as a control. A puncher (5 mm in diameter) was sterilized; Mycelial plugs (5 × 5 mm) were cut from the periphery of 3-day-old colonies of each isolate a mycelia-carrying disc was taken at the edge of the fungal colonies, and the mycelial disc was transferred to a plate with an inoculation needle, with the mycelia facing downward. Four replicates were included for each treatment. Plates were placed in a 25°C biochemical incubator for 4 days, and the colony diameter (minus the original diameter of the inoculation plug) was determined as the average of two perpendicular measurements. Calculate the mycelial growth inhibition rate and a virulence regression equation was established to obtain the half-maximal effective concentration (EC_50_) value. The experiment was performed twice.

### Safety test

The safety test was conducted by referring to “Crop safety evaluation criteria for farm chemicals” and “Indoor test methods for crop safety evaluation of seed treatment agents” NY/T1965.3-2013(People’s Republic of China Agricultural Industry Standard), and the experimental setup was as follows: Before seed sowing, fully developed corn seeds of uniform size were selected for disinfection and placed in sterilized river sand (60 to 70 mesh) in germination boxes(ABS material, transparent, 360mm× 29mm× 12mm in volume) with the moisture content controlled at 60% to 80%. For each treatment, 1 kg of seed was dressed uniformly and air dried. The thifluzamide (24% FS) dosages were set as 192 g a.i. 100 kg^-1^ seed, 96 g a.i. 100 kg^-1^seed, 48 g a.i. 100 kg^-1^ seed, 24 g a.i. 100 kg^-1^ seed, 12 g a.i. 100 kg^-1^ seed, 6 g a.i. 100 kg^-1^ seed, and a control (CK). Thus, a total of 7 treatments were included with 4 replicates per treatment and 50 seeds per replicate. A label was pasted on the side of each germination box with the sample number, species name, and time. Germination boxes were maintained in a GXZ light incubator (28°C, 14 h of light). On the 7th day after establishment, the germination rate, seedling height, root length, root number, and fresh plant weight were measured, and the germination index and vigour index were calculated. The experiment was performed three times.

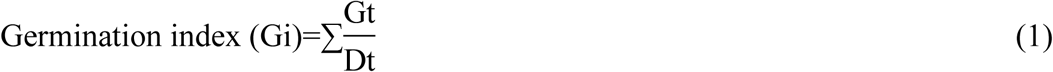

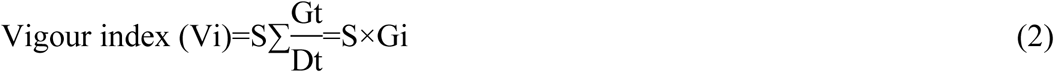

Note: where Gt is the number of germinated seedlings on the T^th^ day; Dt is the corresponding days needed for germination; and S is the fresh weight per plant on the 7^th^ day.

### Greenhouse pot test

The greenhouse pot test included a total of 6 treatments: the 24% thifluzamide (FS) dosages of 48 g a.i. 100 kg^-1^ seed, 24 g a.i. 100 kg^-1^ seed, 12 g a.i. 100 kg^-1^ seed, 6 g a.i. 100 kg^-1^ seed, the control agent tebuconazole at a dosage of 12 g a.i. 100 kg^-1^ seed, and CK. The root activity and chlorophyll content of corn were sampled at the 3-leaf stage. The root activity was determined by the TTC reduction method [23], and the chlorophyll concentration was determined by the extraction method of Ming et al [24,25]. The experiment was performed three times.

### Field fungicide test

The test site was established in Ningyang County of Tai’an City in field plots where the incidence of sheath blight was severe. The test plots had a total acreage of 1,000 m^2^. The soil was loam with uniform fertility, and the irrigation conditions were good. In the 2016 test, seed sowing occurred on June 21, and harvest occurred on September 24; in the 2017 test, seed sowing occurred on June 19, and harvest occurred on September 21. Seeding with mealie socket seeder(Zhengzhou Minle Agricultural Machinery Co., Ltd.), first adjust the sowing depth to 30 mm, insert the tip of the mealie socket seeder into the soil, the seeds fall into the soil, pull out the mealie socket seeder, and level the soil with the foot. Sowing was implemented using the single-seed dibble seeding method with 2 rows per film and plant spacing of 22 cm and row spacing of 45 cm. The dosages of 24% thifluzamide (FS) included 48 g a.i. 100 kg^-1^ seed, 24 g a.i. 100 kg^-1^ seed, and 12 g a.i. 100 kg^-1^ seed; the control fungicide tebuconazole was applied at a dosage of 12 g a.i. 100 kg^-1^ seed; and seed dressing treatments without thifluzamide were taken as a control. Thus, there was a total of 5 treatments in a randomized block design with 3 replicates per treatment, and each plot was 30 m^2^. Corn seedlings were evaluated as follows. One week after planting, 5 sites were sampled in each plot, and 30 plants were surveyed at each site. On the 10^th^ day after sowing, 5 sites were sampled in each plot, and 15 plants were excavated to investigate plant height, stem thickness, root length, and the number of fibrous roots. The fresh plants were weighed, and the root-to-crown ratio was calculated. Before the corn was harvested, 5 sites were sampled for each plot, and samples were brought back to the laboratory for investigation, which included ear length, ear thickness, number of rows per ear, number of grains per ear, and the hundred-grain weight. The yield per 667m^2^ and yield increase rate were calculated as well. The condition index of banded leaf sheath blight in corn was investigated at the small bell stage, large bell stage, tasseling and pollen-shedding stage, silking stage, milk-ripening stage, and wax-ripening stage. At each plot, 5 sites were diagonally sampled, and 20 plants were surveyed at each site to determine the number of diseased plants and the disease grades. The disease rate, condition index, and control effect were calculated according to Eqs. (6), (7), and (8), respectively. The disease grading was conducted according to the grading standards of the International Maize and Wheat Improvement Center (CIMMYT) (Table 1).

**Table 1.** Grading standard for corn sheath blight.

| Disease grade | Typical value | Grading standard |
| --- | --- | --- |
| 0 | 0 | No disease incidence in the whole plant |
| 1 | 1 | Disease incidence at sheaths, at and above the 4<sup>th</sup> sheath below the ear position |
| 2 | 3 | Disease incidence at sheaths, at and above the 3<sup>rd</sup> sheath below the ear position |
| 3 | 5 | Disease incidence at sheaths, at and above the 2<sup>nd</sup> sheath below the ear position |
| 4 | 7 | Disease incidence at sheaths, at and above the 1<sup>st</sup> sheath below the ear position |
| 5 | 9 | Disease incidence at the ear position and at sheaths above the ear position |

### Data processing

All data were processed using SAS statistical software package (version 9.2; SAS). The EC_50_ values of each isolate were calculated by plotting the relative inhibition against the log10 of the fungicide concentration used. To detect differences between treatments, the means of control efficacy were arcsine transformed, then compared with Fisher’s Least Significant Difference test (LSD, P<0.05).

## Results

### Establishment of baseline sensitivity of *Rhizoctonia solani* to thifluzamide in corn

The sensitivity of 102 strains of *R. solani* in corn to thifluzamide was determined using the mycelial growth rate method. It was shown that *R. solani* was highly sensitive to thifluzamide, with an EC_50_ range of 0.0103-0.1942 and an EC_50_ average value of 0.086±0.004 μg/m. The skewness=0.298, kurt=-0.298, and p=0.0884>0.05, which agrees with continuous skewed normal distribution, and the sensitivity frequency distribution had a continuous unimodal curve (Figure 1) and can be used as the baseline sensitivity of *R. solani* in corn to thifluzamide in the Shandong region.

**Fig 1.**
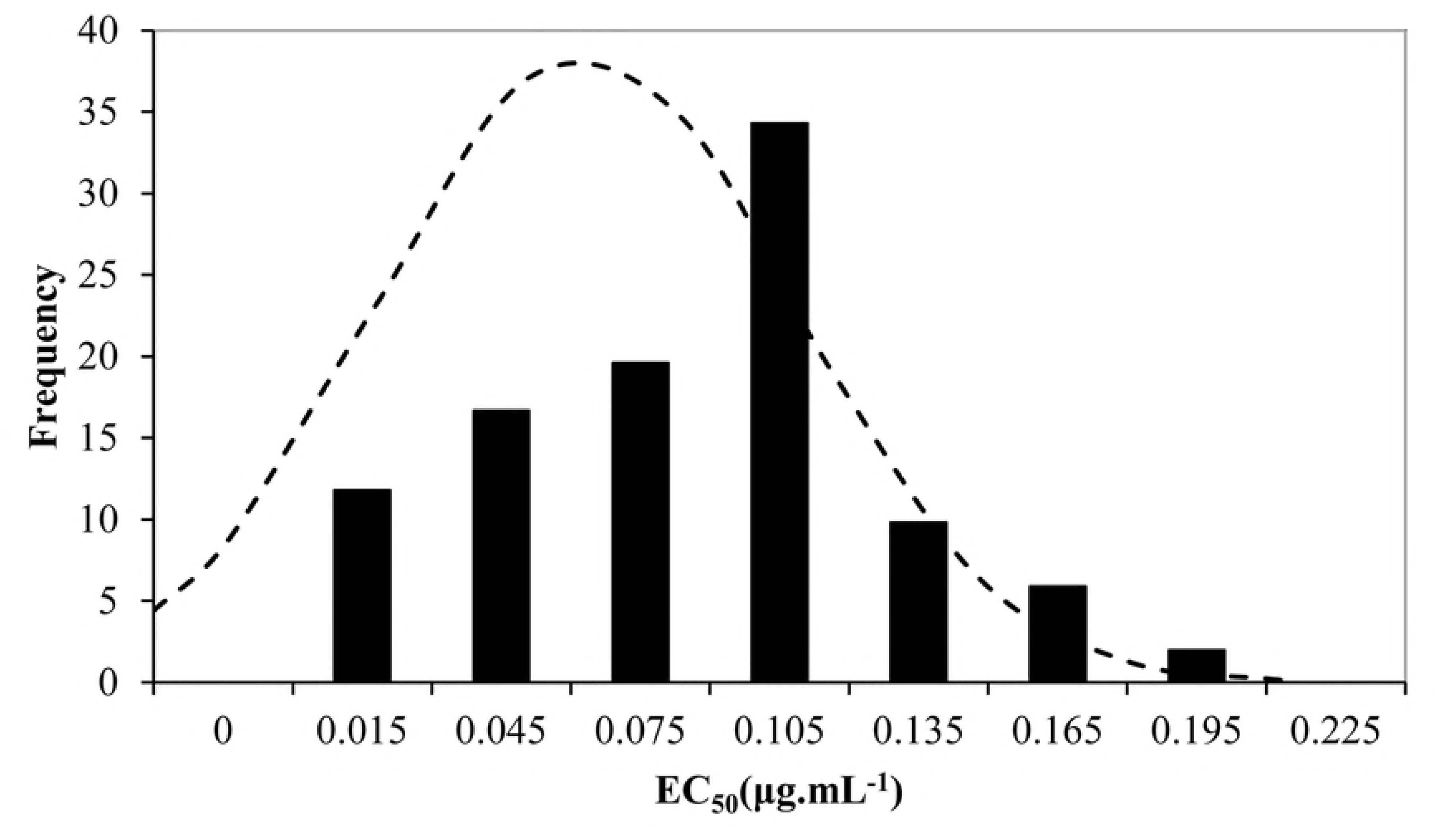
Frequency distributions of 50% effective concentration (EC_50_) of 102 *R. solani* in corn isolates treated with thifluzamide based on mycelial growth. EC_50_ values were calculated by performing a regression of the percentage relative growth against the log_10_ fungicide concentration.

### Safety of thifluzamide in corn

Thifluzamide (24% FS) was generally safe for corn, but excessive use (192 g a.i. 100 kg^-1^ seed) had an adverse effect on indicators, including seedling height, root length, and germination rate. When the dosage was 6-96 g a.i. 100 kg^-1^ seed, corn was safe, and the dosage of 12 g a.i. 100 kg^-1^ seed promoted plant height, root length, root number, the root-to-crown ratio, and the germination index. The dosage of 6 g a.i. 100 kg^-1^ seed had the most favourable effect on the seedling emergence rate, plant fresh weight, and vigour index (Table 2).

**Table 2.** Safety of thifluzamide in corn ^a^.

| Dosage <sup>b</sup><br>(g a.i. 100 kg<sup>-1</sup><br>seed) | Plant height<br>(cm) | Root length<br>(cm) | Root<br>number<br>(piece) | Fresh mass<br>(g) | Germination<br>rate<br>(%) | Shoot ratio<br>(%) | Germination<br>index<br>(%) | Vigour<br>index<br>(%) |
| --- | --- | --- | --- | --- | --- | --- | --- | --- |
| 192 | 7.99±1.05d <sup>c</sup> | 7.15±0.46c | 5.04±0.21a | 1.15±0.05d | 66.67±1.26f | 1.54±0.04d | 9.73±0.08f | 11.19±0.14g |
| 96 | 14.76±0.76bc | 11.73±0.01a | 5.14±0.05a | 1.67±0.17b | 90.00±1.65d | 1.67±0.06c | 13.63±0.09d | 22.71±0.09e |
| 48 | 13.49±0.45bc | 13.03±0.93a | 5.22±0.71a | 1.69±0.20b | 93.33±1.44c | 1.95±0.15ab | 14.30±0.07c | 24.13±0.20d |
| 24 | 14.49±0.01bc | 13.54±0.65a | 5.25±0.32a | 1.71±0.17b | 95.00±2.10b | 1.94±0.04b | 14.88±0.02b | 25.41±0.08c |
| 12 | 17.98±0.02a | 13.81±0.25a | 5.33±0.23a | 1.76±0.07ab | 98.33±1.04a | 2.06±0.05a | 15.58±0.01a | 27.37±0.07b |
| 6 | 15.52±0.26b | 13.42±0.64a | 5.23±0.15a | 1.83±0.04a | 98.33±1.28a | 1.88±0.10b | 15.50±0.01a | 28.39±0.07a |
| CK | 13.07±0.16c | 9.1±0.24b | 4.41±0.28b | 1.45±0.07c | 76.67±1.37e | 1.73±0.12c | 12.33±0.02e | 17.88±0.21f |
| *p*-value | <0.01 | <0.01 | <0.01 | <0.01 | <0.01 | <0.01 | <0.01 | <0.01 |

<sup>a</sup> The experiments performed in the laboratory in 2016.
<sup>b</sup> “Dosage” means the effective concentration.
<sup>c</sup> Mean values followed by the same letter in the columns were not significantly different according to Fisher’s LSD test at *P*=0.05.

### Effects of thifluzamide on root activity and chlorophyll content

Seed dressing with thifluzamide could improve the root activity and increase the chlorophyll content of corn seedlings, among which the dosages of 24 g a.i. 100 kg^-1^ seed and 12 g a.i. 100 kg^-1^ seed had the most significant promotional effect and outperformed the tebuconazole treatment (Figures 2 and 3).

**Fig 2.**
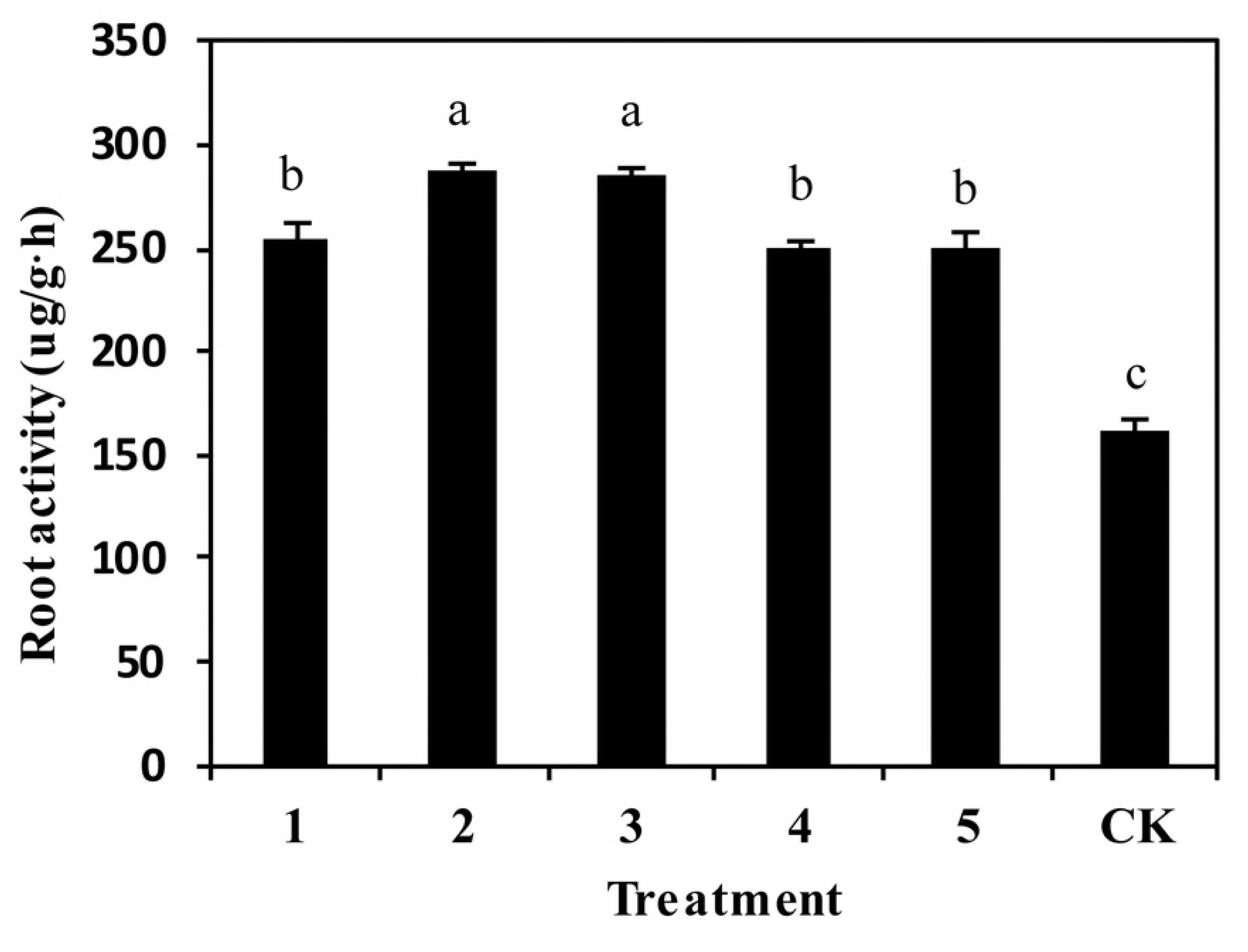
Effect of seed dressing with thifluzamide on the root activity of corn seedlings.

**Fig 3.**
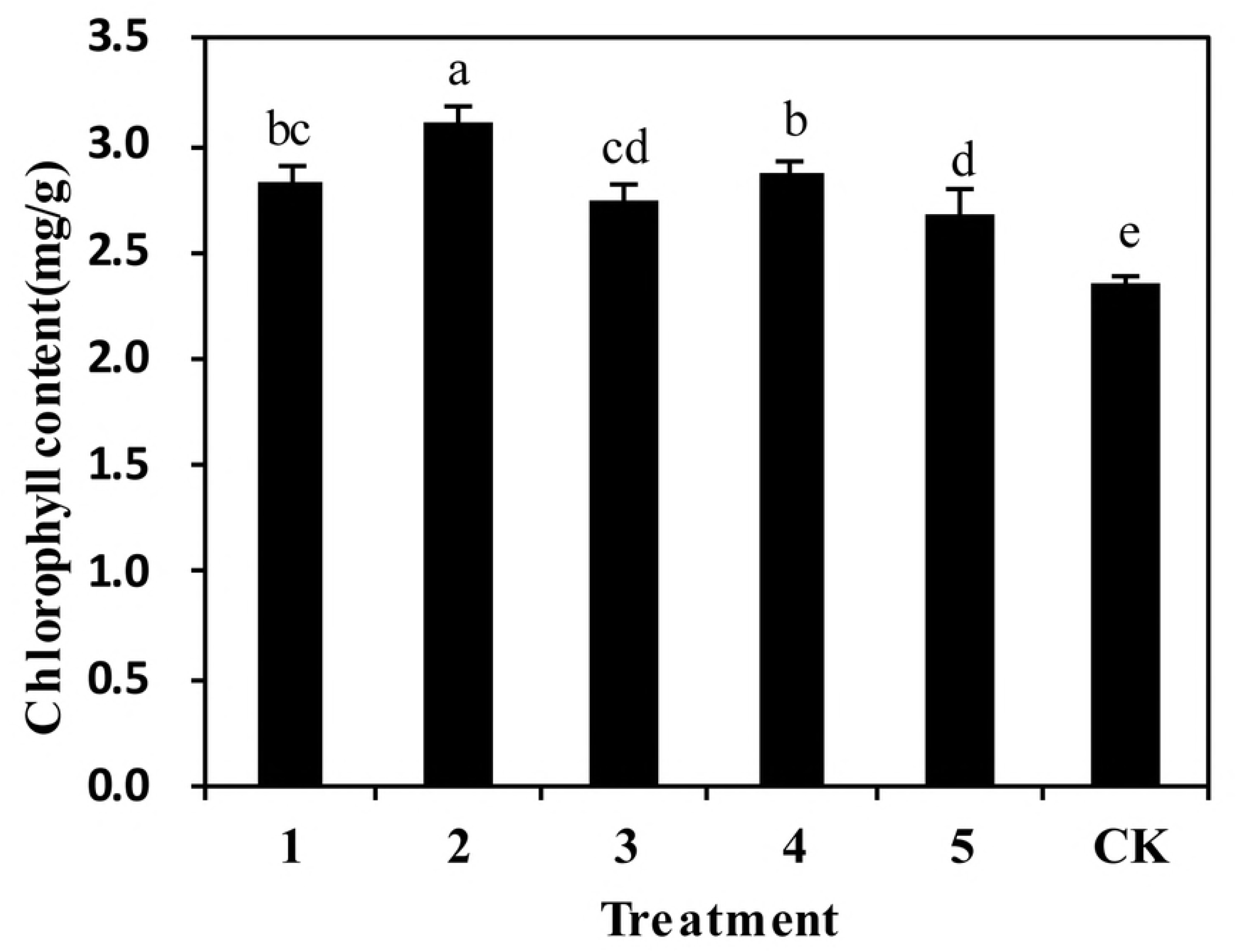
Effect of seed dressing with thifluzamide on the chlorophyll content of corn seedlings.

### Effect of thifluzamide on field emergence of corn

Three dosages of 24% thifluzamide (FS) increased the emergence rate and seedling growth of corn to varying extents. Among them, in 2016 and 2017, the 24 g a.i. 100 kg^-1^ seed dosage had the most favourable effect on the seedling emergence rate, plant height, main root length, fibrous root number, and plant fresh weight. In 2016, The seedling emergence rate was 15.91% higher than the control, and the plant height, main root length, fibrous root number, and plant fresh weight were increased by 4.16 cm, 2.94 cm, 0.87, and 0.64 g, respectively. The dosage of 12 g a.i. 100 kg^-1^ seed had a better promotional effect on stem thickness, which was 0.75 mm higher than that of the control (Table 3). Three doses of thifluzamide (FS) significantly increased the corn root-to-crown ratio, which was obviously better than that under the tebuconazole treatment. Similarly, the 2017 study further validated the 2016 conclusion. 3 dosages of 24% thifluzamide (FS) increased the emergence rate and seedling growth of corn to varying extents (Table 4).

**Table 3.** Effect of thifluzamide on field emergence of corn (2016) ^a^.

| Fungicide | Dosage<br>(g a.i. 100 kg<sup>-1</sup> seed) | Emergence<br>rate<br>(%) | Plant<br>height<br>(cm) | Stem<br>thickness<br>(mm) | Main root<br>length<br>(cm) | Fibrous root<br>number<br>(piece) | Fresh<br>weight<br>(g) | Shoot<br>ratio |
| --- | --- | --- | --- | --- | --- | --- | --- | --- |
| Thifluzamide<br>(FS) 24% | 48 | 92.67±1.16a | 17.01±0.05c | 2.93±0.11b | 14.45±0.15b | 3.47±0.21c | 2.11±0.15bc | 0.57±0.02ab |
|  | 24 | 95.56±0.84a | 18.61±0.08a | 3.10±0.12ab | 16.20±0.07a | 3.89±0.09a | 2.53±0.20a | 0.61±0.02a |
|  | 12 | 93.78±1.86a | 18.06±0.05b | 3.16±0.11a | 15.58±0.16a | 3.76±0.04ab | 2.31±0.23ab | 0.55±0.02b |
| Tebuconazole<br>(FS) 60 g/liter | 12 | 95.11±1.65a | 17.01±0.13c | 2.95±0.03ab | 15.39±0.08a | 3.56±0.15bc | 2.32±0.12ab | 0.54±0.02b |
| CK | - | 82.44±1.92b | 14.45±0.12d | 2.41±0.09c | 13.26±0.35c | 3.02±0.13d | 1.89±0.13c | 0.39±0.02c |
| *p*-value | - | <0.01 | <0.01 | <0.01 | <0.01 | <0.01 | <0.01 | 0.0293 |

**Table 4.** Effect of thifluzamide on field emergence of corn (2017) ^b^.

| Fungicide | Dosage <sup>c</sup><br>(g a.i. 100 kg<sup>-1</sup> seed) | Emergence<br>rate<br>(%) | Plant<br>height<br>(cm) | Stem<br>thickness<br>(mm) | Main root<br>length<br>(cm) | Fibrous root<br>number<br>(piece) | Fresh<br>weight<br>(g) | Shoot<br>ratio |
| --- | --- | --- | --- | --- | --- | --- | --- | --- |
| Thifluzamide<br>(FS) 24% | 48 | 99.33±0.33a <sup>d</sup> | 16.70±0.21c | 2.92±0.04c | 14.24±0.00d | 3.44±0.17b | 2.29±0.08ab | 0.58±0.03a |
|  | 24 | 97.33±0.67a | 19.08±0.24a | 3.17±0.03a | 16.13±0.04a | 4.38±0.25a | 2.69±0.08a | 0.62±0.02a |
|  | 12 | 92.67±0.88bc | 18.09±0.34ab | 3.13±0.03ab | 14.79±0.14c | 3.78±0.20ab | 2.39±0.09ab | 0.55±0.01a |
| Tebuconazole<br>(FS) 60 g/liter | 12 | 98.67±0.33a | 17.36±0.54bc | 2.99±0.01bc | 15.57±0.14b | 3.84±0.25ab | 2.66±0.09a | 0.60±0.01a |
| CK | - | 90.00±0.58c | 14.80±0.13d | 2.49±0.00d | 13.85±0.02d | 3.16±0.10b | 2.19±0.07b | 0.42±0.02b |
| *p*-value | - | <0.01 | <0.01 | <0.01 | <0.01 | <0.01 | 0.0123 | <0.01 |

<sup>a</sup> The experiments performed in the field in 2016. <sup>b</sup> The experiments performed in the field in 2017.
<sup>c</sup> “Dosage” means the effective concentration. <sup>d</sup> Mean values followed by the same letter in the columns were not significantly different according to Fisher’s LSD test at *P*=0.05.

### Effects of thifluzamide on corn yield

Three doses of thifluzamide could increase the ear length, ear thickness, number of rows per ear, and number of grains per ear in the field test of this study. The laboratory seed investigation showed that thifluzamide (FS) could significantly increase the 100-grain weight of corn and the yield per plot. The 24 g a.i. 100 kg^-1^ seed treatment increased the 100-grain weight by 12.47% (2016) and 13.44% (2017) compared with the control, leading to a yield increase of 15.72% (2016) and 14.11% (2017) (Table 5 and 6).

**Table 5.**
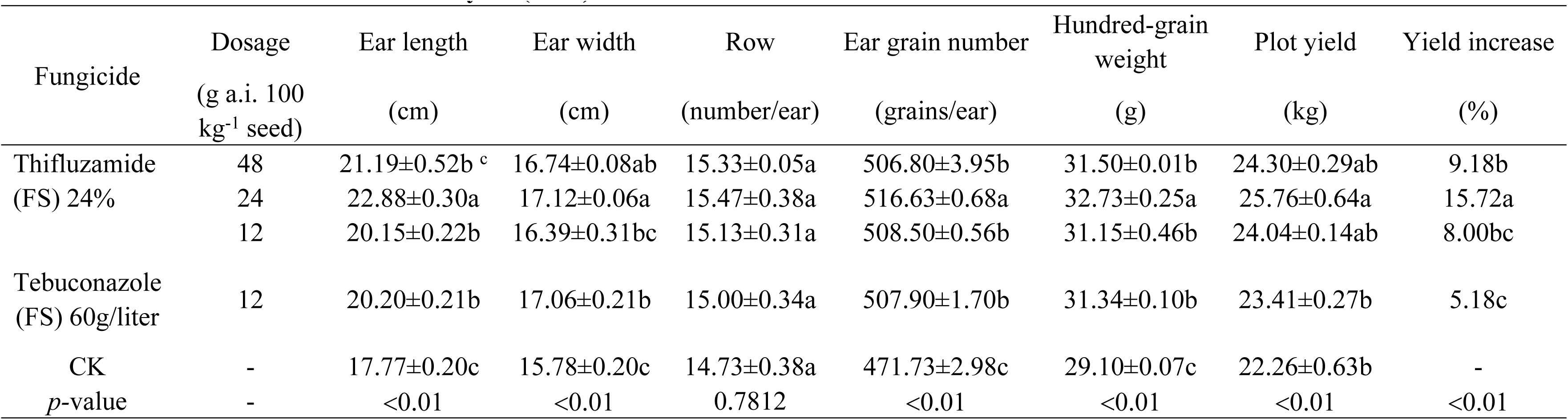
Effects of thifluzamide on corn yield(2016) ^a^.

**Table 6.**
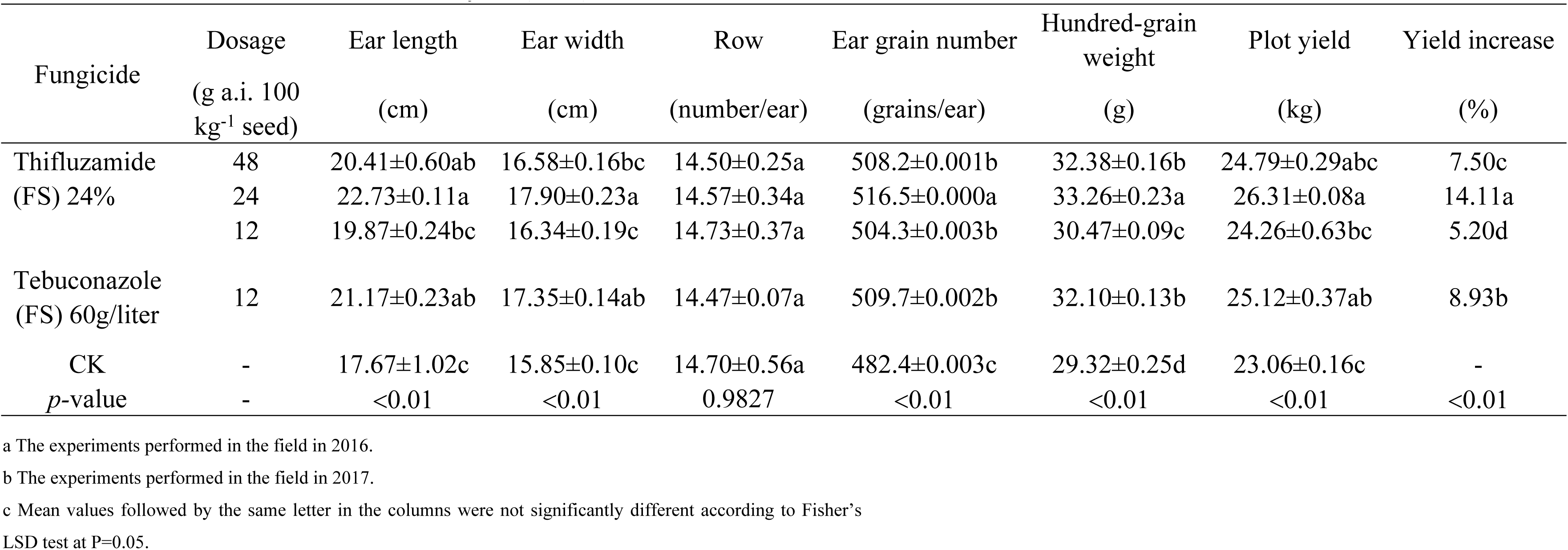
Effects of thifluzamide on corn yield(2017) ^b^.

### Effects of thifluzamide on the prevention of banded leaf sheath blight in corn in the field

In the field test of this study we found that there were fewer incidences of banded leaf sheath blight in corn from the seedling stage to the large bell stage, during which the control effect was high. The tasseling and pollen-shedding stage was the disease-spreading period, with high temperature and humidity conditions being conducive to the spread of sheath blight, and the maturity stage was the abrupt surge period of the disease. The 2-year field trial showed that 3 doses of thifluzamide (FS) had good control effects on banded leaf sheath blight in corn throughout the entire growth period and significantly reduced the incidence of banded leaf sheath blight in corn during the high-incidence period. Among these, the dosage of 24 g a.i. 100 kg^-1^ seed had the optimal field control effect, and the control effects during the small bell stage, large bell stage, tasseling and pollen-shedding stage, silking stage, milk-ripening stage, and wax-ripening stage were 100%, 66.73%, 52.8%, 67.81%, 68.48%, and 62.68% (2016), respectively, and 74.97%, 63.17%, 50.90%, 53.60%, 61.42%, and 55.88% (2017). Through field observation and data analysis, the disease rate in the plots under the seed dressing with thifluzamide treatment was significantly higher during the period from the late wax-ripening stage to corn harvest than during other stages (Table 7).

**Table 7.**
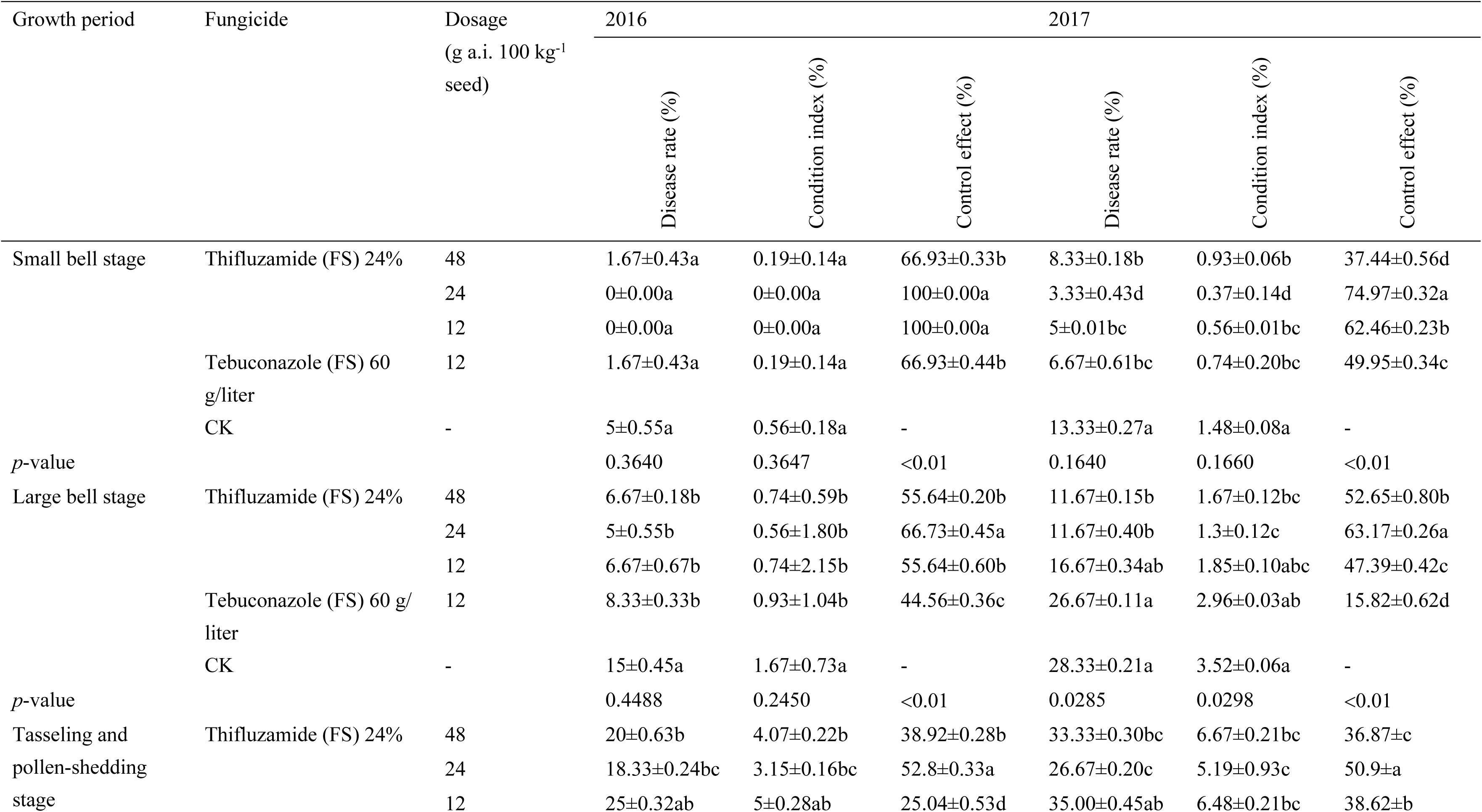
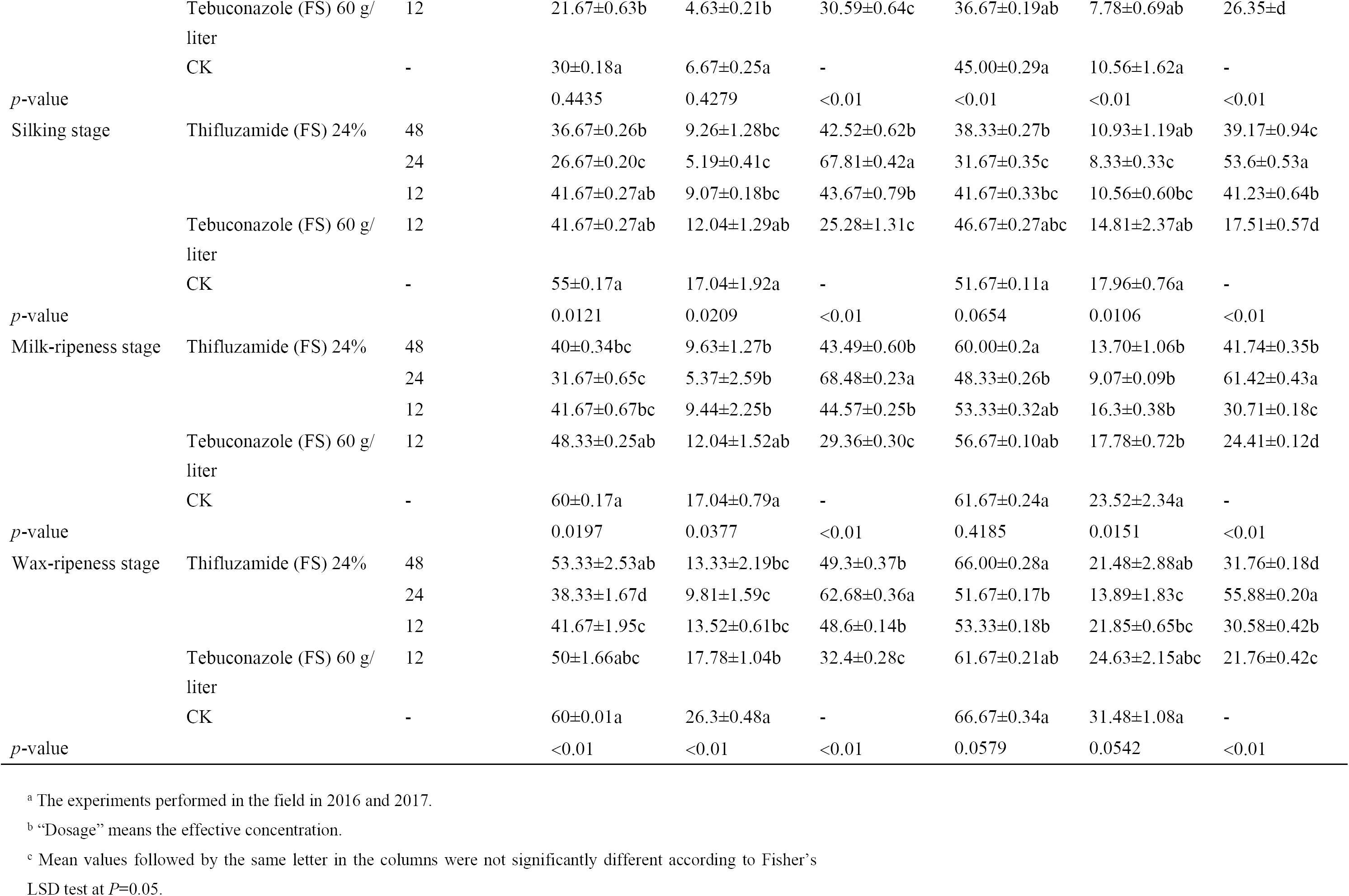
Effects of thifluzamide on the prevention of corn sheath blight in the field in 2016 and 2017 ^a^.

## Discussion

Being a fungicide of the succinate dehydrogenase inhibitor (SDHI) type, thifluzamide inhibits the synthesis of succinate dehydrogenase [26], thereby preventing pathogens from transmitting electrons in the mitochondria [27], thus inhibiting their growth [28]. Studies have shown that thifluzamide has high inhibitory activity against *R. solani* and can be used as a more effective substitute for boscalid and Jinggang mycin to control sheath blight [29]. Hence, we established the baseline sensitivity of *R. solani* in corn to thifluzamide and found that it was highly sensitive. Of the 55 fungicides listed by the Fungicide Resistance Action Committee (FRAC), the SDHI class is growing at the fastest rate among the new compounds that have been produced and put on the market [26]. As an SDHI fungicide, thifluzamide has high biological activity, but it only has a single action site, so it runs a high risk of drug resistance [30]. A previous study found that the risk of resistance to thifluzamide is moderate in *R. solani*, which can develop resistance to QoI fungicides, and the Fungicide Resistance Action Committee (FRAC) states that the use of this fungicide should be in accordance with the manufacturer’s recommended effective dose, with particular attention to adhering to safety intervals [31]. In this study, we did not spray and reduced the number of fungicide applications, and the optimal dosage was determined in the indoor safety test and the greenhouse pot experiment using the seed dressing method. When the thifluzamide dosage (24% FS) was 6-96 g a.i. 100 kg^-1^ seed, seed coating with this fungicide was safe for corn. The field study found that the seed coating treatment at the dosage of 24 g a.i. 100 kg^-1^ had the highest field control effect on banded leaf sheath blight in corn and could provide a theoretical basis for control using thifluzamide. Thifluzamide has strong adsorption capacity in the soil, but its adsorption intensity is weak, with 19.5%-54.0% digestion in 90 days [32]. In the field test of this study, the disease rate of banded leaf sheath blight in corn at each plot treated with thifluzamide (FS) was found to significantly increase after the late milk-ripening stage, but the control effect was still higher than that of the blank control and the control fungicide. It can be basically guaranteed that thifluzamide would not be applied to corn during the whole growth period.

Currently, the main prevention and control measures for banded leaf sheath blight in corn are chemical. Jiang stated that the control of banded leaf sheath blight in corn should be based on agricultural methods, with seed treatment with chemical agents being the main approach. The study by Xue et al. showed that the control effect of banded leaf sheath blight in corn was significantly different when fungicide application occurred during different growth stages, and the jointing stage was the best period for application [33]. Taking the traditional fungicide Jinggang mycin as an example, although 2 consecutive applications by leaf sheath spraying in the early tasseling stage has a good control effect, the application method is time consuming, labourious, and causes severe air pollution at large dosages that is unsafe for natural enemies, humans, and livestock, which has caused the chemical to be banned in many countries. In addition, spraying is ineffective for controlling soil-borne diseases and has a short duration of effectiveness. Furthermore, multiple applications are required, and the awareness of disease control is weak among farmers. Therefore, it is necessary to develop efficient, safe and time-saving fungicides. In this study, the control effect of thifluzamide suspension (FS) on banded leaf sheath blight in corn in the field was significantly higher than that under seed dressing with the control fungicide tebuconazole. Compared with traditional fungicide agents and fungicide application methods, thifluzamide has the advantages of an increased utilization rate, guaranteeing precise application, reduced application frequency, which saves seeds and fungicide, and reduced production costs, and it has broad prospects for development. In conjunction with the call of the public for environmental protection, biological control has also made great breakthroughs in recent years. Chaurasia et al. isolated antagonized *Bacillus subtilis*, which produces diffusive and volatile compounds that can induce the separation of the tested mycelia and conidia [34], and Stein found that the peptide and non-peptide metabolites produced by *B. subtilis* have antibacterial activities [35]. However, the effectiveness of biological control is greatly affected by environmental conditions, and it is difficult to meet expectations. In a greenhouse test, the effect of biocontrol with *B. subtilis* was lower than that of Jinggang mycin [36]; meanwhile, the control effect of *Trichoderma* spp. against banded leaf sheath blight in corn can reach up to 68.52% [37]. Considering various aspects such as economic benefits and natural environmental conditions, biological control still needs to be developed. Many studies have shown that SDHI fungicides have good health protection effects on plants and can promote crop growth and enhance the ability of crops to tolerate adverse environments. A previous study by Lde and Dubois showed that Benodanil can prevent and control diseases caused by *Rhizoctonia* in a variety of crops and can increase yield [38], and field trials have found that Carboxin can stimulate wheat growth and increase yield [32]. When thifluzamide is applied at 240 g/L, rice leaves become broader, thicker, and greener, and rice stalks exhibit enhanced toughness, which promotes robust growth. Worthing CR et al. found that compound products such as penflufen, Emesto, and EverGol can improve the crop viability, improve resistance in plants, and increase crop quality [39]. Through a greenhouse pot test in this study, the effects of seed coating using a thifluzamide suspension agent on the root activity and chlorophyll content of corn were preliminarily determined, which showed that the fungicide had a significant promotional effect and has further research value.

## Supporting Information

S1 Table. Meteorological data sheet during the test (2016)

S2 Table. Meteorological data sheet during the test (2017)

## Acknowledgements

This study has received funding from the Technology Research and Demonstration on Reduction of Chemical Fertilizers and Pesticides in Summer Maize in Huang-Huai-Hai Region (SQ2018YFD020062-4), the Provincial Major Technological Innovation Program of Agricultural Application in Shandong, and the Shandong “double first-class” award (SYL2017-XTTD11).

## Author Contributions

Conceived and designed the experiments: DLS XYJ. Performed the experiments: DLS CTY FLH XS SSS. Analysed the data: DLS HLT. Contributed reagents/materials/analysis tools: DLS XDL JWZ. Wrote the paper: DLS.

## References

1. Akhtar J, Jha VK, Kumar A, Lal HC. Occurrence of banded leaf and sheath blight of maize in Jharkhand with reference to diversity in Rhizoctonia solani. Asian Journal of Agricultural Sciences. 2009;1(2):32–35.

2. Rashed OZA, Abdullah SNA, Alsultan WMK, Ahmad K. Genetic variability of Rhizoctonia spp. isolated from different hosts and locations. 2017.

3. Baker R, Martinson CA. Epidemiology of diseases caused by Rhizoctonia solani. Rhizoctonia Solani Biology & Pathology. 1970.

4. Pascual, Toda, Raymondo, Hyakumachi. Characterization by conventional techniques and PCR of Rhizoctonia solani isolates causing banded leaf sheath blight in maize. Plant Pathology. 2010;49(1):108–118.

5. Pascual CB, Raymundo AD, Hyakumachi M. Efficacy of hypovirulent binucleate Rhizoctonia sp. to control banded leaf and sheath blight in corn. Journal of General Plant Pathology. 2000;66(1):95–102.

6. Hirrel MC. First Report of Sheath Blight (Rhizoctonia solani) on Field Corn in Arkansas. Plant Disease. 1988;72(7).

7. Zhao M, Zhang Z, Li W, Pan G. Advances on research of banded leaf and sheath blight of maize. Plant Protection. 2006;32(1):5–8.

8. Jhm S, Salazar O, Rubio V, Keijer J. Identification of Rhizoctonia solani associated with field-grown tulips using ITS rDNA polymorphism and pectic zymograms. European Journal of Plant Pathology. 1997;103(7):607–622.

9. Ogoshi A. Ecology and Pathogenicity of Anastomosis and Intraspecific Groups of Rhizoctonia Solani Kuhn. Annual Review of Phytopathology. 1987;25(1):125–143.

10. Sneh B, Burpee L, Ogoshi A. Identification of Rhizoctonia species. Brittonia. 1991.

11. Li HR, Wu BC, Yan SQ. Aetiology of Rhizoctonia in sheath blight of maize in Sichuan. Plant Pathology. 1998;47(1):16–21.

12. Abendroth L, Elmore RW, Boyer M, Marlay S. Corn growth and development. 2011.

13. Jackson TA. Reemergence of Goss’s Wilt and Blight of Corn to the Central High Plains. Plant Health Progress. 2007.

14. Yan JM, Zheng J, Hua-Zhi YE, Zhang M, Qin Y. Damage and Yield Loss in Corn Caused by Corn Sheath Blight. Journal of Maize Sciences. 2008;16(5):123–125.

15. Tagele SB, Sang WK, Lee HG, Kim HS, Lee YS. Effectiveness of multi-trait Burkholderia contaminans KNU17BI1 in growth promotion and management of banded leaf and sheath blight in maize seedling. Microbiological Research. 2018.

16. Li S. Experiment on the control of maize sheath blight by triadimefon. 2003.

17. Mu W, Wang Z, Bi Y, Ni X, Hou Y, Zhang S, et al Sensitivity determination and resistance risk assessment of Rhizoctonia solani to SDHI fungicide thifluzamide. Annals of Applied Biology. 2017;170(2):240–250.

18. Wei-Qun HU, Song HM, Zhu WG, Zhang RR, Chen J. Synergistic and Field Effects of Thifluzamide and Fludioxonil against Rhizoctonia solani. Agrochemicals. 2014.

19. Arias Rivas B, Mcgee D, Burris JS. Evaluación del potencial de polímeros como agentes envolventes de fungicidas en el tratamiento de semillas de maíz para el control de Pythium spp. Maria Dengosa. 1998;67(2):152.

20. Kunkur VK, Hunje R, Biradarpatil NK, Vyakarnahal BS. Effect of Seed Coating with Polymer, Fungicide and Insecticide on Seed Quality in Cotton During Storage. Karnataka Journal of Agricultural Sciences. 2010;20(1).

21. Pereira CE, Oliveira JA. Qualidade fisiológica de sementes de milho tratadas associadas a polímeros durante o armazenamento Performance of corn seeds treated with furazin and maxin in association with polimers, during storage. Ciência E Agrotecnologia. 2005;29(6):1201–1208.

22. Avelar, GoncalvesSousa SA, Defiss FV, GuilhermeBaudet, LeopoldoPeske, Teichert S. The use of film coating on the performance of treated corn seed. Revista Brasileira De Sementes. 2012;34(2):186–192.

23. Baozhang B, Jin J, Huang L, Song B. Improvement of TTC Method Determining Root Activity in Corn. Maizeence. 1994.

24. Arnon DI. COPPER ENZYMES IN ISOLATED CHLOROPLASTS. POLYPHENOLOXIDASE IN BETA VULGARIS. Plant Physiology. 1949;24(1):1–15.

25. Ming H, Chun-Sheng HU, Zhang YM, Cheng YS. Improved Extraction Methods of Chlorophyll from Maize. Journal of Maize Sciences. 2007.

26. Sierotzki H, Scalliet G. A review of current knowledge of resistance aspects for the next-generation succinate dehydrogenase inhibitor fungicides. Phytopathology. 2013;103(9):880–887.

27. Sun H, Wang C, Li W, Zhang A, Deng Y, Chen H. Characterization of Rhizoctonia cerealis sensitivity to thifluzamide in China. Crop Protection. 2015;69:65–69.

28. He L, Cui K, Ma D, Shen R, Huang X, Jiang J, et al Activity, translocation and persistence of isopyrazam for controlling cucumber powdery mildew. Plant Disease. 2017;101(7).

29. Chen Y, Zhang AF, Wang WX, Zhang Y, Gao TC. Baseline sensitivity and efficacy of thifluzamide in Rhizoctonia solani. Annals of Applied Biology. 2012;161(3):247–254.

30. Ajayi-Oyetunde OO, Butts-Wilmsmeyer CJ, Bradley C. Sensitivity of Rhizoctonia solani to succinate dehydrogenase inhibitor and demethylation inhibitor fungicides. Plant Disease. 2016;101(3).

31. Liang-Kong LI, Yuan SK, Pan HY, Wang Y. Progress in Research on SDHIs Fungicides and Its Resistance. Agrochemicals. 2011.

32. Gupta S, Gajbhiye VT. Adsorption-desorption, persistence and leaching behavior of thifluzamide in alluvial soil. Chemosphere. 2004;57(6):471–480.

33. Xue T, Fu J, Zhou R. The epidemiology of corn sheath blight and its preventive treatment. Journal of Maize Sciences. 2008;16(1):126–128.

34. Chaurasia B, Pandey A, Palni LM, Trivedi P, Kumar B, Colvin N. Diffusible and volatile compounds produced by an antagonistic Bacillus subtilis strain cause structural deformations in pathogenic fungi in vitro. Microbiological Research. 2005;160(1):75–81.

35. Stein T. Bacillus subtilis antibiotics: structures, syntheses and specific functions. Molecular Microbiology. 2005;56(4):845–857.

36. Mao T, Ye H, Yuhua Q. Study on Biocontrol of Maize Sheath Blight(Rhizoctonia solani) with Bacillus subtilis Strain. Chinese Agricultural Science Bulletin. 2016.

37. Zhang G, Wen C. Biocontrol of maize sheath blight with {\sl Trichoderma} spp. Journal of Plant Protection. 2005;32.

38. Lde LDB, Dubois C. Distribution of thiabendazole-resistant Colletotrichum musae isolates from Guadeloupe banana plantations. Plant Disease. 2011;81(12):1378–1383.

39. Worthing CR, Walker SB. The pesticide manual, a world compendium. British Crop Protection Council. 1991; (2):148–148.

